# The spatial extent of a single lipid’s influence on bilayer mechanics

**DOI:** 10.1101/2021.01.20.427479

**Authors:** Kayla C. Sapp, Alexander J. Sodt, Andrew H. Beaven

## Abstract

To what spatial extent does a single lipid affect the mechanical properties of the membrane that surrounds it? The lipid composition of a membrane determines its mechanical properties. The shapes available to the membrane depend on its compositional material properties, and therefore, the lipid environment. Because each individual lipid species’ chemistry is different, it is important to know its range of influence on membrane mechanical properties. This is defined herein as the lipid’s mechanical extent. Here, a lipid’s mechanical extent is determined by quantifying lipid redistribution and the average curvature that lipid species experience on fluctuating membrane surfaces. A surprising finding is that, unlike unsaturated lipids, saturated lipids have a complicated, non-local effect on the surrounding surface, with the interaction strength maximal at a finite length-scale. The methodology provides the means to substantially enrich curvature-energy models of membrane structures, quantifying what was previously only conjecture.

## INTRODUCTION

Amphiphilic lipids form the foundation of the bilayers that function as a physical barrier, surrounding and protecting the living cell. Their collective mechanics determine the rates of many critical biological processes, including both viral entry and exit, as well as how the cell recycles membrane signaling proteins as a fundamental element of regulating its response to stimuli [1]. Biological membranes are composed of hundreds of chemically distinct lipid species [2], each with an individual effect on surface mechanics [3–7]. Yet the spatial form of that effect, and how it depends on the physical interactions between lipids, is largely unknown.

Interactions between lipids (either individually or collectively) determine the stable shapes and patterns [8, 9] that critically influence biological processes. For example, biological membranes sit close to a two-dimensional compositional phase transition [10, 11] and experiments suggest they tune their lipid composition to maintain that proximity [12]. Material properties also determine the shapes a biological membrane can support. Intermediates in membrane fusion and fission are prime examples of curved structures that are favored by certain lipids. To determine the energy of a membrane shape, and how lipids affect its stability, the curvature (*c*) at a point on the shape is compared to the *spontaneous curvature* (*c*_0_) of the lipids at that point. A commonly employed elastic energy functional, the Helfrich/Canham (HC) Hamiltonian [13, 14] is proportional to the squared deviation of *c* and *c*_0_.

This model is able to describe a number of biologically relevant phenomena. The relative stability of flat and highly curved hexagonal phases is determined by lipid headgroup and acyl chain chemistry [6, 15, 16]. Adding positively-curved lysolipids blocks fusion while negatively-curved arachidonic acid favors fusion [17]. A lipid’s *c*_0_ determines how it segregates between the leaflets of sonicated vesicles [18, 19]. These experimental observations can be explained applying the HC model, which, as typically applied, generally assumes heterogeneous lipid compositions. This assumption is frequently well justified. However, it is an open question as to how single lipids influence local material properties.

For example, consider a simple assumption (analyzed in more depth below): that a lipid determines bilayer mechanical parameters within a “footprint” of area *A_p_* around it (i.e., a local effect). Here, the subscript *p* refers to treating an individual lipid as a particle diffusing in the membrane surface. The local HC energy for the lipid is:

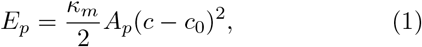

where *κ_m_* is the bending modulus of the leaflet. Within this model and assumptions, a force acts to move the lipid to where *c* is equal to *c*_0_. Using common values for the elastic constants (*κ*_m_ = 7.7 kcal/mol, *A* = 65 Å^2^, *c*_0_ = [−29 Å]^−1^) applicable to high-curvature-favoring phosphatidylethanolamine (PE) lipids [4], yields *E_p_* = 0.30 kcal/mol on a flat surface, or a Boltzmann weight of 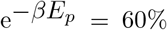 compared to a small, highly curved surface with curvature *c*_0_. For larger structures, enrichment is even more modest and challenging to detect as a single-lipid effect [20].

An alternative assumption is that the effect on spontaneous curvature is spread over the entire bilayer (i.e., a longer-range effect). In this case, there is no force driving enrichment based on spontaneous curvature. We define the spatial range of a single lipid’s influence on a leaflet’s mechanical parameters as the “mechanical extent;” (ME) it is placed in mathematical terms below as a spatial weight, *w*(*x, y*), which is in turn extracted from molecular simulation data.

The difference between local mechanical extent and longer-range mechanical extent is critical for determining how a heterogeneous lipid environment [21], be it transient or persistent, will support a membrane reshaping process like endocytosis or viral budding. For local extent, the influence of a lipid is independent of the frequency *q* of the bilayer undulation; a concentration of curvature sensitive lipids will strongly promote shapes even at small length-scales. For well-distributed extent, only low *q* modes will feel the influence of the lipid. A third possibility, which we present evidence for below in the case of saturated lipids, is that a lipid’s influence is maximal at a particular value of *q*. Discussed below, this physical mechanism favors modulation of the shape of laterally-separated lipid phases, potentially explaining how lipid composition can tune a length-scale in quaternary mixtures [22]. Previously, a finite length-scale was predicted to emerge only as a result of the interplay between tension and compositional curvature-coupling [23–26].

Ternary systems of dioleoyl (DO)-phosphatidylcholine (PC, or DOPC), DOPE, and DO-phosphatidyl serine (PS, or DOPS) are used to test the redistribution behavior of known lipids. Of the simple lipids of the plasma membrane (with two acyl chains and no carbohydrate units), those with PE headgroups have the most significant curvature, *c*_0_ = [−29 Å]^−1^ [3, 4], and simulations reproduce the curvature of PE accurately [27]. Extending the simulations to saturated lipids (palmitoyl sphingomyelin, i.e., PSM and dipalmitoyl-PC, i.e., DPPC), mixed with palmitoyl-oleoyl-PC (POPC) tests the effect of acyl chain on ME. Generally, results indicate a lipid’s ME is localized. However, an interesting effect emerges for the saturated lipids as their fraction increases; simulations indicate a finite length-scale of curvature preference on the order of nanometers. As discussed below, this has novel implications for why complex mixtures including saturated lipids give rise to rich patterning as part of liquid-liquid lateral phase separation.

## THEORY AND METHODS

### Continuum energetics and the definition of mechanical extent

The following sections establish two curvature-related quantities necessary to determine the mechanical extent of a lipid’s influence on bilayer mechanics. The first is the *transverse curvature bias*, 〈*c*_**q**_〉(*z*) (see Eq. 16), defined as the apparent curvature sampled by a lipidic surface when the positions of the constituent lipids are measured at a height *z* above the bilayer midplane. This quantity is necessary to correct for systematic bias determining the local curvature around a lipid that is introduced when choosing an internal coordinate for a lipid’s lateral position.

The unbiased position is called the *neutral surface* [28, 29] and is determined by where in a lipid the transverse curvature bias is zero. Once we demonstrate how to compute curvature free from the bias of the lipid coordinate system, we can define the *spontaneous curvature spectrum*, Δ*c*_0_(**q**) (see Eq. 28). The quantity Δ*c*_0_(**q**) is the difference in spontaneous curvature of a lipid and the surface background implied by the dynamic redistribution of the lipid on an undulation with wavevector **q**. We demonstrate how the **q** dependence of Δ*c*_0_(**q**) determines the mechanical extent of the lipid, as we define it.

#### 1. Preliminary Fourier description of lipid surfaces

In this work, out-of-plane undulations, *h*(*x, y*), and ME, *w*(*x, y*), will be described in Fourier space:

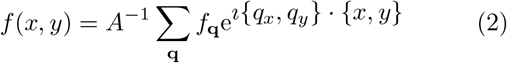

where the coefficients, *f*_**q**_, are determined by

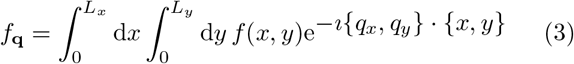

and where the function *f* can be *h* or *w*. For the Fourier transform of the periodic bilayer, Eq. 2, only *q*-space wavevectors compatible with the periodic boundary conditions are non-zero, that is:

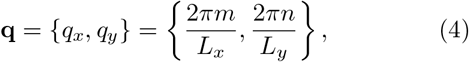

where *m* and *n* are integers and *L_x_* and *L_y_* are the periodic cell dimensions. Since these functions are realvalued, the property 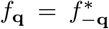 must hold. Therefore, only half of the modes are independent and a set of wave-vectors can be defined using the shorthand “ {**q** > 0} ” as [30]:

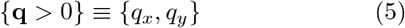

such that

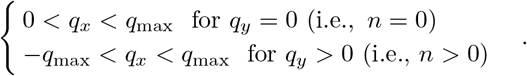

Note that negative *q_y_* are only included implicitly in the set through 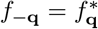. This contains all the necessary information for the independent modes and allows for the treatment of the real and imaginary components of the Fourier amplitudes separately. Therefore, computations can be done on a “per mode” or “per degree of freedom” basis. Moving forward we use the “per mode” formalism; however we do comment on the differences when necessary.

##### Fluctuations in planar membrane curvature

Consider a single undulating mode of a bilayer described with Fourier amplitudes as in Eq. 3. Computing the HC Hamiltonian energy density,

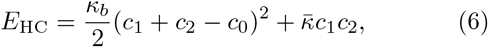

where *κ_b_* (the bilayer bending rigidity) and 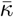 (the saddlesplay modulus) require expressions for the principal curvatures *c*_1_ and *c*_2_ in terms of *h*_**q**_. The principal curvatures are quadratic coefficients describing the parabolic variation of the surface away from the tangent plane. Straight-forward formulas are available for computing them for general surfaces [31]. For nearly planar simulations with small fluctuations, the use of the *linearized Monge gauge* simplifies mathematical analysis. The Monge gauge parameterization begins with

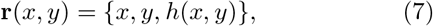

and continues by dropping terms of higher order than 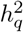 when computing observables. This truncation is applicable only to a small deformation, which includes most thermally induced undulations at physiological temperatures. The principal curvatures are now

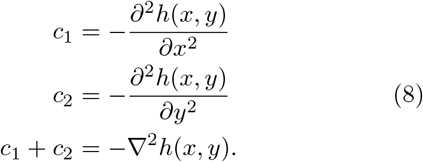

The leading signs are chosen so that curvature is positive if the upper (+*z*) leaflet headgroups are outside of a convex surface. The energy as a function of *h*_**q**_ is

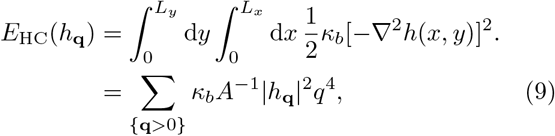

with *A* = *L_x_L_y_*. The surface normal to first order in *h_q_* is

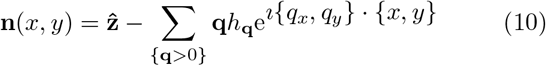

where 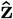 is the unit *z* direction normal to the flat membrane. Auxiliary surfaces can then be defined that are displaced along the membrane normal, **n**, a distance *z*:

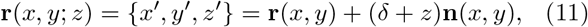

where *δ* is the so-called *neutral surface* at which curvature and area-compression energetics are uncoupled [28, 32], and at which the surface pivots at nearly constant area. This definition will be used subsequently to describe variations in curvature at different *transverse* positions along the bilayer normal. Notationally *z* is used as a parameter, with each value defining a distinct surface (displaced by *z* along the normal) with distinct curvature.

According to Eq. 9, the modes **q** do not couple energetically, therefore the partition function *Z*_**q**_ for a single mode can be expressed as:

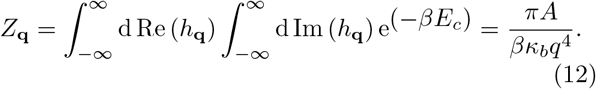

where the integral is taking over the real and imaginary parts of a single Fourier amplitude, *h*_**q**_. The whole system partition function ℤ for the independent modes is the product of the individual functions:

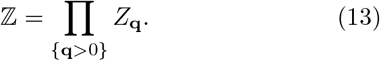

##### The transverse curvature bias 〈*c*_q_〉(*z*)

Embedded in the linearized Monge gauge is the assumption that the lipids are evenly distributed on the bilayer surface [33]. Under this assumption, the average curvature of the surface (zero) is also the average curvature experienced by a lipid (for a single component bilayer). However, this assumption is only valid if a lipid’s position is measured at *δ* (notationally defined here as *z* = 0, see Eq. 11). For example, in the case of negative curvature, lipids will appear to be more concentrated above the neutral surface as a consequence of this systematic biased sampling of their position. The lipid normal tends to point toward negative curvature. This leads to systematic bias of the sampled curvature in terms of which internal coordinate of the lipid is used to track the lipid. The top panel of Figure 1 depicts the origin of the bias graphically. The assumption of a uniform distribution of lipids will be broken.

**FIG. 1:**
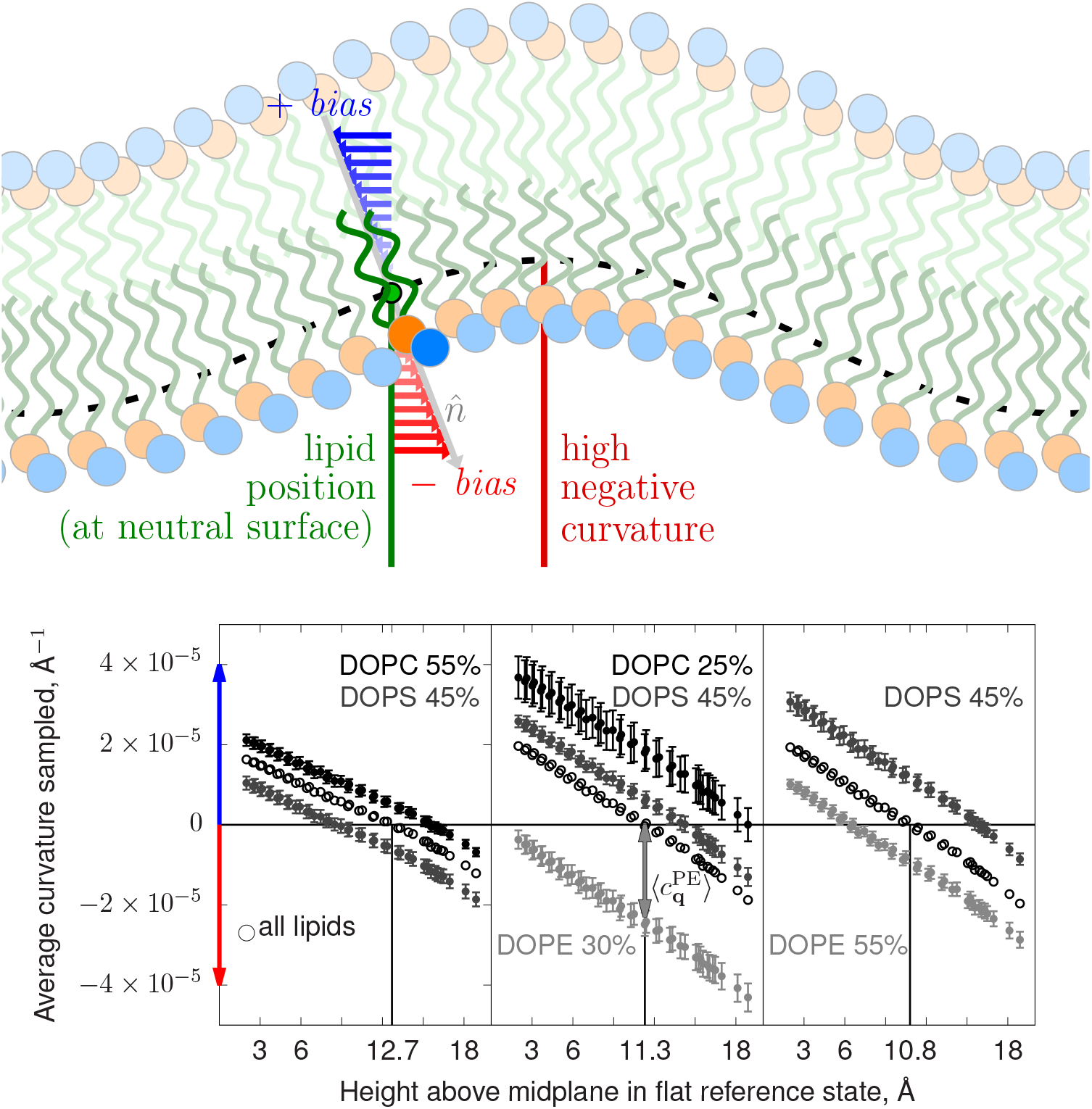
Top: A cartoon illustrating the curvature-sampling bias created by tracking a lipid’s position away from the neutral surface. Colored arrows indicate how measuring a lipid’s position off the neutral surface (dashed black line) biases either toward negative curvature when sampled too close to the head groups (red arrows) or toward positive curvature when sampled too far into the acyl chain region (blue arrows). Bottom: Transverse curvature bias profiles, 〈*c*_**q**_〉 (*z*), for mixtures including DOPC, DOPE, and DOPS. Profiles including all lipids are shown in black. The average curvature per lipid (vertical axis) is reported as a function of the atom used to measure a lipid’s lateral position. The atom’s identity is mapped to its height in a nearly planar initial state (horizontal axis). The lipid’s average curvature is recorded at the neutral surface determined using the average over all lipids. The arrow for DOPE in the center panel indicates the shift due to lipid spontaneous curvature. The red and blue colors indicate the sign of curvature according to the cartoon above. Note that the data points for each lipid’s profile are not independent; they are correlated by being spatially fixed by the molecular geometry. Error bars are standard errors of the mean.

Transverse curvature bias only applies to molecular motions that are correlated with collective undulation; the magnitudes of tilt (i.e., orientational noise distributed around the normal) and protrusion (lipids sliding along the normal) will not influence the observed curvature when averaged over sufficient time.

Without loss of generality, consider a mode with height variation along only the *x* direction (*q_y_* = 0). For an arbitrary atom *i* in a lipid, its location will be measured at position **r**(*x*; *z*), where *z* is the average displacement of atom *i* from *δ*. As previously mentioned, the density of the lipid will not be uniform (in general) when measured by the position of atom *i*. Instead, a spatially varying metric factor

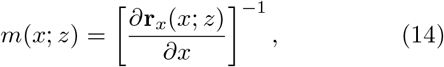

again parameterized by *z* and with **r**_*x*_(*x, z*) as the first component of the vector defined in Eq. 11, quantifies the change in density. Here the ratio of a change in **r**_*x*_ to a change in *x* is the relative area of the displaced surface at *z* to the base surface. The inverse relationship of area to density determines the inverse proportionality in Eq. 14. While this appears to define a full *three-dimensional* model of the leaflet similarly to Ref. [34], the goal is to determine the proper definition of the *two-dimensional* surface model where curvature should be measured. Three-dimensional mechanical properties can be determined by tracking the deformation of a leaflet’s atoms, for example, to determine Poisson’s ratio [35].

The expected curvature sampled by an atomic site *i* for some mode **q** = {*q_x_,* 0}, integrated over the patch area, is

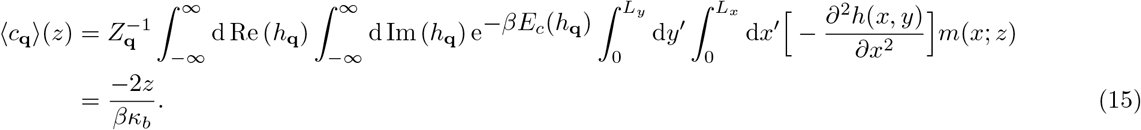

Where, consistent with the linearized Monge gauge, terms of order 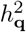 and higher have been dropped. Per unit area, the curvature will thus be

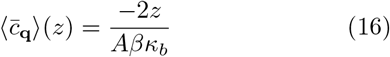

where 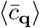 is the curvature per mode (*per area*, as denoted by the bar). Note that there is no expected dependence on *q* for the curvature per mode; the subscript serves to indicate that this is a “per mode” value. This is the definition of the transverse curvature bias.

Per *degree-of-freedom*, the value is halved; that is, there are two degrees of freedom denoted by **q**. Note that *δ* in Eq. 7 does not appear when limiting the expansion to 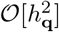. The logic above dictates that the reference surface should have a uniform distribution of lipids as it is deformed. In practice, some distribution *ρ*(*x*) of lipid position will be sampled at atom *j*. This distribution can be written as *ρ*(*x*) = *ρ*_0_*m*(*x*; *z*′ − *δ*), where the position of the neutral surface *δ* may be unknown (here *z*′ = *z* + *δ*). By plotting 〈*c*_**q**_〉(*z*′) against *z*′, *δ* can be determined by finding where 〈*c*_**q**_〉(*z*′) is zero. This is the basis for the transverse curvature bias profiles plotted below in Figure 1.

##### *The spontaneous curvature spectrum,* Δ*c*_0_(**q**) *of a single lipid*

Consider an individual lipid with spontaneous curvature *c*_0*,i*_ in a bilayer composed of lipids with spontaneous curvature *c*_0_. We model the ME of a lipid by a function *w*(*x, y*) that is the fractional impact of the lipid as a distance from the internal coordinate chosen by analysis of the bias above. The HC energy is then modified by a change Δ*E_p_* based on the lipid’s parameters:

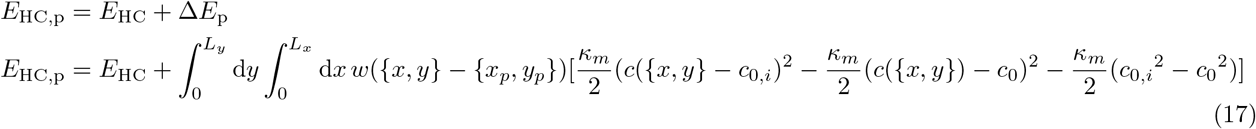

where *w*(*x, y*) is normalized such that

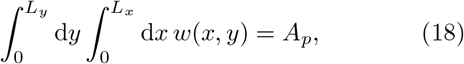

where *A* is the area of the surface integrated over, and *A_p_* is the area of the single lipid. Note that a constant term, 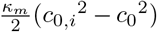 has been subtracted that would influence only the chemical potential.

To compute the partition function now requires integrating over the surface of the membrane while including the contribution from the particle at {*x_p_, y_p_*}:

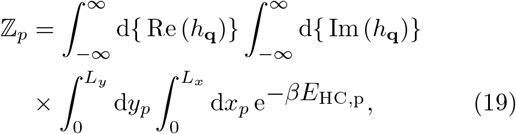

where here *E*_HC,p_ depends on the Fourier amplitudes ({*h*_**q**_}) and the lipid position (*x_p_*, *y_p_*). The integral is taken over the single particle position and the set of *h*_**q**_ Fourier coefficients. As before, the membrane modes are separable, therefore we can write the partition function for a single mode and single particle as:

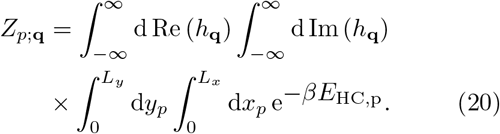

In this model, lipids are not directly coupled to each other, and so only couple through membrane fluctuations and so would be similarly separable in the case of multiple particles.

Inserting the Fourier series representation of *w* into Eq. 17 yields

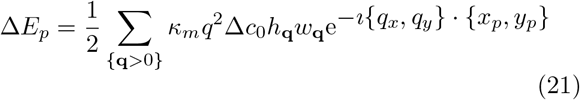

where Δ*c*_0_ = *c*_0*,i*_ − *c*_0_. The expectation value for curvature is then

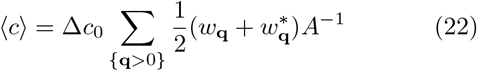

The average curvature sampled depends on the wavelength of the undulation based on *w*(*x, y*).

Now consider the simplest case of point-wise (local) ME,

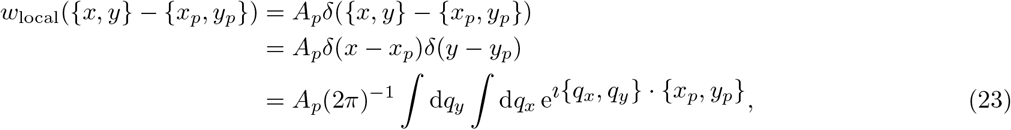

that is,

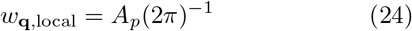

The single-particle, single-mode partition function *Z*_**q**;*p*_, to second order in Δ*c*_0_, is:

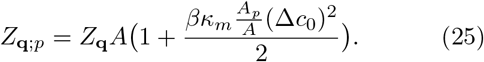

The average curvature sampled by the lipid is

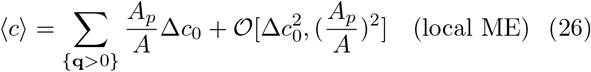

The curvature sampled using a local extent function, *w_local_*, is *q* independent.

The integration steps necessary to arrive at the partition functions and average curvatures are variations on

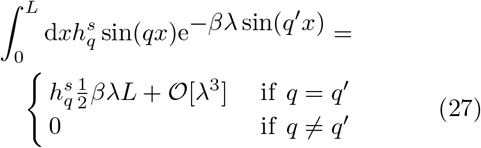

where *q* = 2*πmL*^−1^, *q*′ = 2*πnL*^−1^ with integer *m* and *n*, 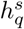 is the magnitude of a sinusoidal undulation, and *λ* is a coupling constant, e.g., Δ*c*_0_. The sin function in the exponential is the weight due to curvature energetics, while the pre-exponential factor is the magnitude of curvature itself.

The ME can be extracted from the spontaneous curvature spectrum:

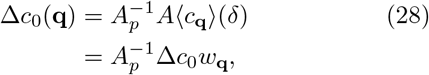

where 〈*c*_**q**_〉 indicates the expectation value of curvature sampled along mode **q**. The simulated area *A* and sampled curvature 〈*c*_**q**_〉 depend on simulation size, whereas the spectrum itself does not. Note that when *w*_**q**_ is included in the definition of Δ*c*_0_(*q*), a constant Δ*c*_0_ enters the formula. This reflects the normalization of *w*_**q**_ in Eq. 18. With this definition, *w*_**q**_ may be negative if, for example, the sign of the spontaneous curvature switches at a particular value of *q*.

##### *The q* = 0 *limit from the lateral pressure profile*

Spontaneous curvature is typically inferred from molecular simulations using the lateral pressure profile (LPP) method [28, 36]. When integrated over a single leaflet the calculation yields the derivative of the leaflet free energy with respect to curvature,

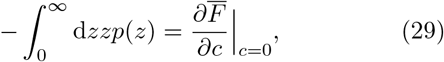

evaluated at zero curvature, similarly to how a virial expression yields the derivative of the free energy with respect to volume, i.e., the pressure. Interpreting the simulated free energy derivative through the HC model of an individual lipid’s effect on bilayer mechanics, Eq. 17, yields:

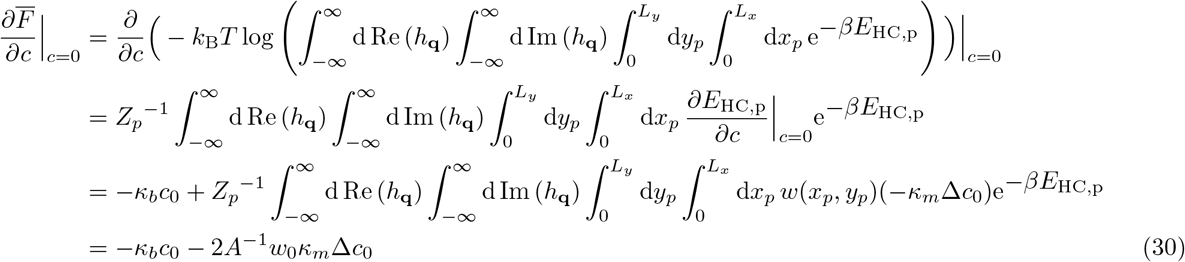

where here we have assumed that the ME has no explicit curvature dependence, which is consistent with the formulation in Eq. 17. The last line follows because all non-zero-frequency contributions to *w*_**q**_ average to zero at net zero curvature:

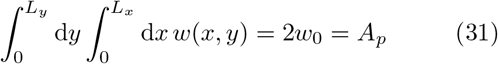

The single point at zero in the spontaneous curvature spectrum is computed from the standard form:

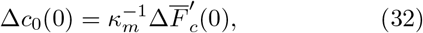

where 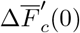 is the difference of 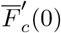 between the lipid under consideration and the average value of the total bi-layer (average values may be weighted by *A_p_* if differences can be estimated).

#### 2. Interpretation of w(x, y)

If *w*(*x, y*) is assumed to be radially symmetric, its Fourier transform can be written as a one dimensional Hankel transformation [37]:

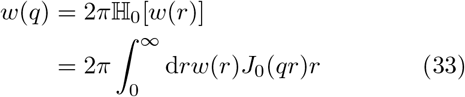

where here *w*(*q*) is similarly symmetric with respect to the orientation of **q** and so only depends on magnitude, and *J*_0_ is a Bessel function of the first kind. The inverse is equivalent, with the exception of the accumulated factors of 2*π*:

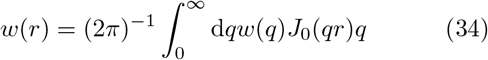

Qualitatively, we interpret the spectrum in terms of three observations. First is how the curvature spectrum attenuates at high *q*. If the spectrum is fit well by an exponential decay, the ME in *q* and real space will be:

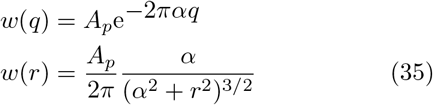

where *α* is the decay coefficient for *q* (see p. 6 of Ref. [38]), this is a Lorentzian. Similarly, if the spectrum more closely matches a Gaussian distribution,

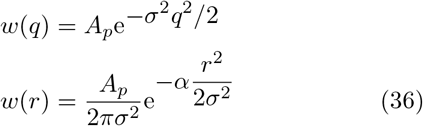

the ME is also a Gaussian. To distinguish between the two is not critical, rather, it is useful to extract a qualitative measure of the ME, such as the Gaussian or Lorentzian width.

Second is whether the spectrum meets the zero-*q* limit established by the LPP, *c*_0*,q*=0_. Large systems are necessary to simulate long modes with slow relaxation times that would inform on the low *q* behavior of *w*(*x, y*). However, if the simulated spontaneous curvature spectrum is inconsistent with *c*_0*,q*=0_, we infer that a component of the spontaneous curvature effect is below the *q* range of the spectrum. That is, the simulation was too small to detect the full ME, or too short to capture the slow relaxation of small *q* modes with reliable statistics.

Third is whether a particular length-scale emerges from the mid-*q* variation of the spontaneous curvature spectrum. This is indicated by a peak or valley in the spectrum, and as discussed further below, would suggest a mechanism for curvature-coupled modulation of the shape and size of lateral compositional inhomogeneity.

## Molecular Dynamics

### Build and simulation parameters

Two sizes of systems were built and simulated: i) large systems with an elongated *y*-axis for computing spontaneous curvature spectra including the effect of low *q* undulations and ii) small, square-patch systems for computing 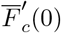 via the LPP. Large-patch systems (1,320 total lipids – 660 lipids / leaflet) were divided into three main groups: i) DOPC:DOPE:DOPS at 55:0:45, 25:30:45, and 0:55:45 mole fractions; PSM:POPC at 10:90, 20:80, and 30:70 mole fractions; and DPPC:POPC at a 20:80 mole fraction. Additionally, small-patch systems (200 total lipids – 100 lipids / leaflet) were simulated: i) 80:0:20 DOPC:DOPE:DOPS; ii) PSM:POPC at 0:100, 5:95, 10:90, 12:88, 15:85, 17:83, 20:80, 30:70, and 40:60 mole fractions. Simulations are referred to by their main lipid constituents in bold face, with DO lipids shortened to their headgroup chemical acronym. For example, DOPC:DOPE:DOPS at 25:30:45 relative composition is **PC**_25_**PE**_30_**PS**_45_.

All systems were built using the CHARMM-GUI *Membrane Builder* protocol [39–43]. Minimization and initial relaxation steps were performed using NAMD [44] as prescribed by CHARMM-GUI. All systems were simulated with a constant temperature of 310.15 K, anisotropic pressure (*x* and *y* coupled; zero surface tension) of 1 bar, and used the CHARMM all-atom force field [45, 46]. Non-bonded interactions were switched off between 10–12 Å, and long-range electrostatics were handled by PME with a spacing of less than 1 Å. All bond lengths involving hydrogen were constrained [47, 48].

### Large-patch simulations with Amber for spontaneous curvature spectra

Following the initial relaxation steps, the large patch systems were converted into AMBER format [49, 50] using ParmEd. The simulations were run using the Amber 18 GPU implementation of PMEMD [51–53]. The temperature was controlled by a Langevin thermostat with a friction coefficient of 1 ps^−1^. Constant pressure was maintained by a Monte Carlo barostat. A 2 fs time step was used with coordinates saved every 200 ps.

### *Small-patch simulations with NAMD for* Δ*c*_0_ *at q* = 0

The small patch systems had constant temperature maintained by a Langevin thermostat with a 1 ps^−1^ damping frequency, and constant pressure was maintained by a Nosé-Hoover Langevin piston [54, 55] with a 50 fs oscillation period and a 25 fs damping time scale. A 1 fs time step was used and coordinates were saved every 200 ps for analysis. LPPs were obtained post-simulation using 250 slabs along *z* using a patched version [56] of NAMD (v2.12).

### *Computation of h*_q_ *and* 〈*c*〉_q_

The instantaneous Fourier spectrum is computed by resolving the height *h*(*x, y*) of the all-atom membrane on a discretized grid of bins, with a maximum spacing of 15 Å (*q*_max_ = 0.21). The number of bins for a particular dimension was computed using ceil(*L/*15 Å). The height is initially computed on a per-leaflet basis. The final height *h_a_* of a grid point labeled by *a* was taken as the mean of the two leaflet heights.

The Fourier amplitude *h*_**q**_ of a particular mode **q** was computed as

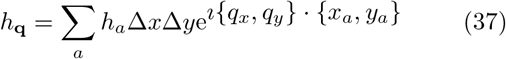

where *h_a_* is the height of grid point *a* with lateral coordinates *x_a_* and *y_a_*, and Δ*x*Δ*y* is the area of the patch that contributes to grid point *a*. As the trajectory was processed to compute *h*_**q**_, the *x* and *y* coordinates of each lipid were also recorded.

The instantaneous curvature at a lipid was then computed as ∇^2^*h*(*x, y*):

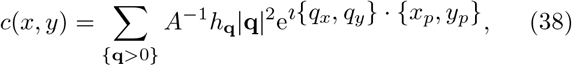

which is real by virtue of *h*(*x, y*) being real as stated above. The *q*-dependent curvature is:

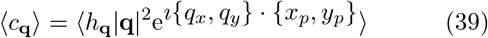

where 〈〉 indicates sampling over the trajectory of the simulation; it depends on the correlation of *h*_**q**_ and the lipid’s position, {*x, y*}.

## I. RESULTS AND DISCUSSION

### The transverse curvature bias determines where to measure lipid position on an undulating surface

Figure 1, at bottom, shows the transverse curvature bias, 〈*c*_**q**_〉 (*z*) computed from Eq. 16, for DOPC, DOPE, and DOPS, including the average over all lipids. The lateral positions of the lipids were measured separately using the heavy atom positions of the forcefield. Each atom has a corresponding Δ*z* computed from the nearly flat initial condition (the first ten nanoseconds, following the standard CHARMM-GUI pre-equilibration sequence). As expected, when measuring the lateral position using atom sites near the acyl chains (bilayer middle) the observed curvature is biased to positive values, while the opposite is true when position is measured with head-group atoms, consistent with the cartoon. The neutral surface, *δ*, where the lateral distribution is uncoupled to curvature, is shown with a vertical line intersecting the labeling of the horizontal axis. This analysis yields the appropriate atom for computing the spontaneous curvature spectrum. Note that the neutral surface atom appears to be *q* dependent, moving closer to the water interface at higher *q*.

The difference in average curvature sampled implies the lipids’ *c*_0_. For a dilute mixture of a lipid like PE in PC or PS, Eq. 16 would provide the difference in spontaneous curvature, Δ*c*_0_, between the target lipid and the background. For PE mixtures like those simulated here, the effect will wane as PE becomes the majority species and is forced to regions of positive curvature simply by self-exclusion. Additionally, mode amplitudes will be enhanced by this weak dynamic redistribution of the lipids to their preferred curvature.

The transverse curvature bias may also be used for robust estimates of the bending modulus; this is described below in the Appendix.

#### A. The *q* = 0 point of the spontaneous curvature spectrum computed from 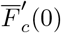

Values of 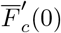 are available for DOPC (0.061 ± 0.0025 kcal/mol/Å) and DOPE (0.2281 ± 0.0036 kcal/mol/Å) in tandem with bending moduli (for both lipids, 17.0 kcal/mol) from Ref. [57]. For DOPS, highly anionic bilayers require a correspondingly large ion concentration for counterbalance. Along with this is the necessity for a sufficiently large water layer to dissipate the layer effects of high salt concentration [58]. Accordingly, the value of 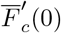 for DOPS was inferred from a relatively low coverage 20% DOPS/20% DOPC mixture.

The value of 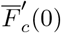 for PSM is determined by fitting the linear coefficient of 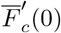 for mixtures of PSM and POPC as the fraction of PSM goes to zero. This is necessary because the value of 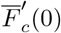 for 100% PSM is not necessarily equal to the contribution of a PSM monomer to the 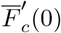 in which PSM is a minority lipid. The correspondence of properties when a lipid is in the minority or majority appears to apply to mixtures of DOPE and DOPC [59]. Note that it is the influence of the *monomer* that is sampled by the curvature spectrum of PSM dilute in POPC. The variation of 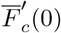 with PSM mole-percentage is plotted in Figure 2. The slope of 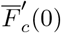 with percentage PSM in POPC is −4 × 10^−4^ ± 5 × 10^−4^ kcal/mol/Å/% when averaged between 0% and 10% and 1.4 10^−3^ 2 10^−4^ kcal/mol/Å/% when averaged between 0% and 20%. Statistically, these data indicate non-linear variation of curvature stress with sphingomyelin content. The full characterization of this non-linear variation, also observed for PSM/DOPE mixtures [59], is beyond the scope of this present work. The fit implies that the contribution to 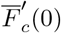 for monomers of PSM is 0.14 0.02 kcal/mol/Å, taking the data from 0% to 20% as the reference.

**FIG. 2:**
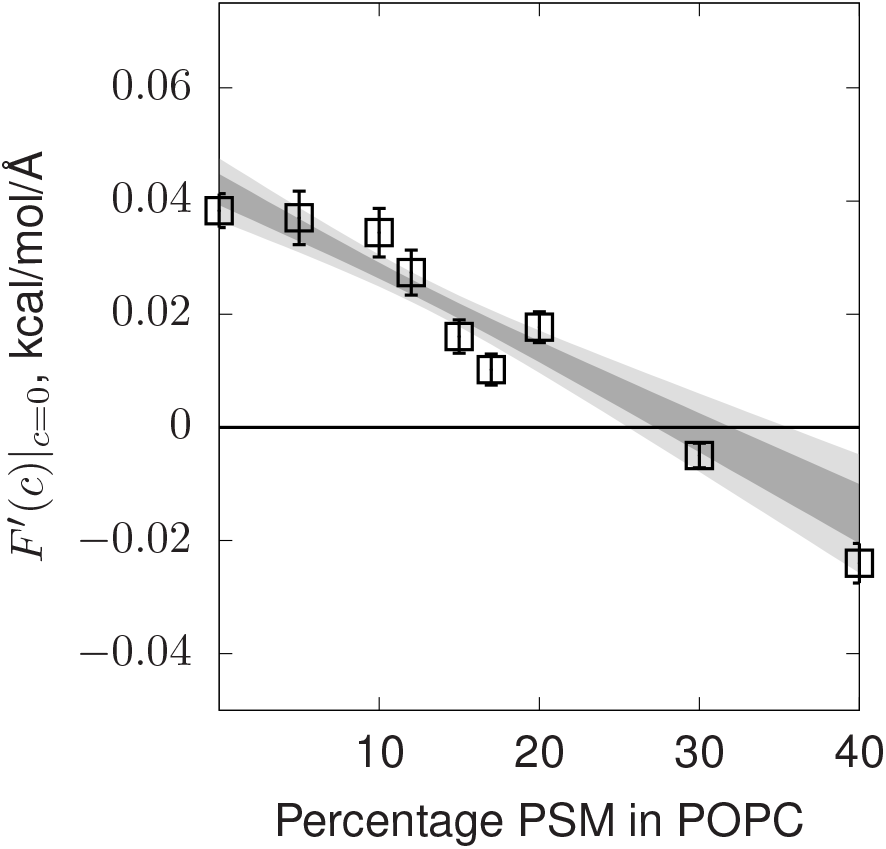
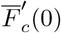 versus percentage of PSM in a POPC bilayer. Error bars are standard errors of the mean. The dark shaded regions are within one standard deviation for a linear fit of their respective data with PSM > 60%. The lighter shaded regions are two standard deviations.

#### B. Fitting *w*(*q*) for a qualitative measure of mechanical extent

Fitting the spontaneous curvature spectra to Eq. 35 yields a model of the spatial range of the individual lipids via the function *w*(*q*). As we have described it, the spontaneous curvature spectrum is a combination of *c*_0_ multiplied by the extent function *w*(*q*). In the extensive experiments performed by Rand and co-workers, e.g. Refs. [3, 4, 6, 7, 16], *c*_0_ is extracted from a model relating osmotic stress to the strain of hexagonally packed lipidic cylindrical monolayers (the inverse hexagonal phase). This yields the zero-frequency contribution to the spectrum.

In this work, the fit is complicated by applying different methodology, (analysis of the LPP) to extract the *q* = 0 point. The LPP has very high statistical precision, but a caveat must be provided; for DOPC, DOPE, DOPS, and DPPC we assume that the parameter *c*_0_ is the same at 100% composition as it is when dilute. If the value of *c*_0_ inferred reflects the lipid matrix, we refer to this as a non-additive contribution to bilayer mechanics. This is clearly an issue for sphingolipids and cholesterol but does not appear to be significant for DOPE and DOPC [59]. The full numerical results of the fits are listed in Table S1 of the supporting material.

##### 1. The q-dependent spontaneous curvature spectra of unsaturated lipids with varying headgroup chemistry indicate primarily localized mechanical extent

The spontaneous curvature spectra for mixtures of DOPC, DOPE, and DOPS are shown in Figure 3. These data are computed from both lipid redistribution information from large simulations (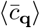, closed points) as well as 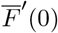 inferred from LPPs (open points at *q* = 0). A lipid with localized extent has a spontaneous curvature spectrum that is constant across *q*, see Eq. 24. The spectra obtained in Figures 3–5 appear to be roughly consistent with an asymptote to a constant value at high *q*, consistent with local extent. It is expected but reassuring that two completely different methodologies (the LPP and dynamic redistribution methods) give consistent Δ*c*_0_. Although qualitatively the methods are consistent at low *q*, there are statistical variations that appear inconsistent with completely localized ME.

**FIG. 3:**
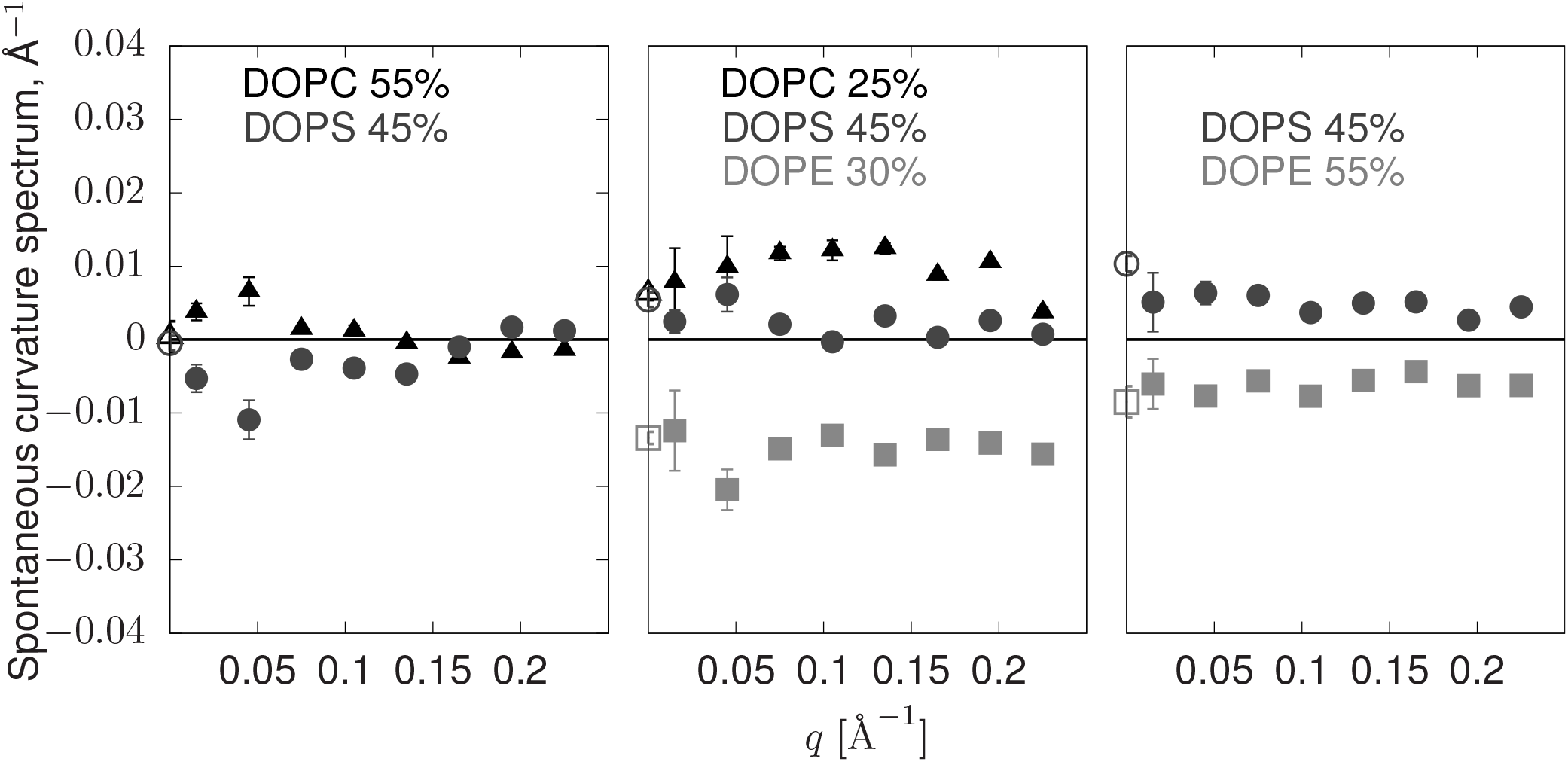
Spontaneous curvature spectra sampled from **PC**_55_**PS**_45_, **PC**_25_**PE**_30_**PS**_45_, and **PE**_55_**PS**_45_. DOPC is shown in black triangles, DOPS in gray circles, and DOPE in lighter gray squares. Open circles at *q* = 0 are computed from the LPP. The data is a histogram of values. Error bars are ± the standard error of the histogram bin mean.

In accord with the observation that the fits are largely local, a robust fit of Eq. 35 can be performed by first subtracting out the value of Δ*c*_0_(*q*) at high *q*, Δ*c*_0_*k*_local_*A_p_*. The magnitude of the residual spectra, *k*_non−local_, indicates the delocalized extent. The ME is fit with:

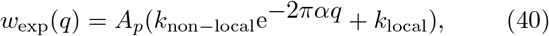

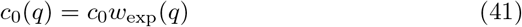

with *c*_0_*k*_non−local_ and *α* as least-squared fit parameters, while *k*_local_ is computed directly from the apparent asymptote above *q* ≥ 0.165. Including all three parameters simultaneously in a least squares fit yields unphysical parameters for very localized spectra, because for small *α* the constant and exponential terms become linearly dependent.

We report here the relative magnitude of the local and non-local pieces with the percent of the effect attributed to the non-local (exponential) factor. In the case where the local and non-local switch signs, e.g., DOPS in **PC**_55_**PS**_45_, absolute values are applied for reporting the percent contribution. Qualitatively, if the spectra are flat across the span of *q* reported from the simulations, they will tend to have large *k*_local_ and thus the same spontaneous curvature for any surface undulation.

There is minimal non-local extent in the **PC**_55_**PS**_45_, **PE**_55_**PS**_45_, and **PC**_25_**PE**_30_**PS**_45_ simulations shown in Figure 3. In both the **PE**_55_**PS**_45_ and **PC**_25_**PE**_30_**PS**_45_ simulations, the introduction of non-local extent either contributes weakly (e.g. *k*_non−local_ = 13% for DOPC and 7% for DOPE in **PC**_25_**PE**_30_**PS**_45_) or does not change the statistical significance of the fit, for DOPE and DOPC. For DOPS in all three of these mixtures, a rather short-ranged *α*, < 3 Å, with magnitude *k*_non−local_ > 60%, improves the fit significantly over a completely local fit. The somewhat small value of *α* is still consistent with the size of a lipid (with an area per lipid ca. 65 Å^2^), although the Lorentzian form of the real-space extent has a long tail. In summary, for these lipids with common headgroup and unsaturated tail chemistry, we find largely localized extent with a small non-local, but still short-ranged, contribution.

##### 2. The complex spontaneous curvature spectra of saturated lipids indicate non-local extent

Figure 4 shows the spontaneous curvature spectra of three PSM/POPC mixtures. The spectrum of PSM drops significantly at high *q* where it displays negative curvature. At low *q*, where collecting statistics is challenging due to the slow relaxation times, error bars are large but generally consistent with an estimate from the LPP that PSM has positive spontaneous curvature. A key feature of increasing PSM in the simulation is the emergence of positive curvature between *q* = 0.05 and *q* = 0.2 Å^−1^. These points are indicated in Figure 4 with black outlines. As apparent outliers, if the highlighted points were not used in the fit their deviation would be even more striking.

**FIG. 4:**
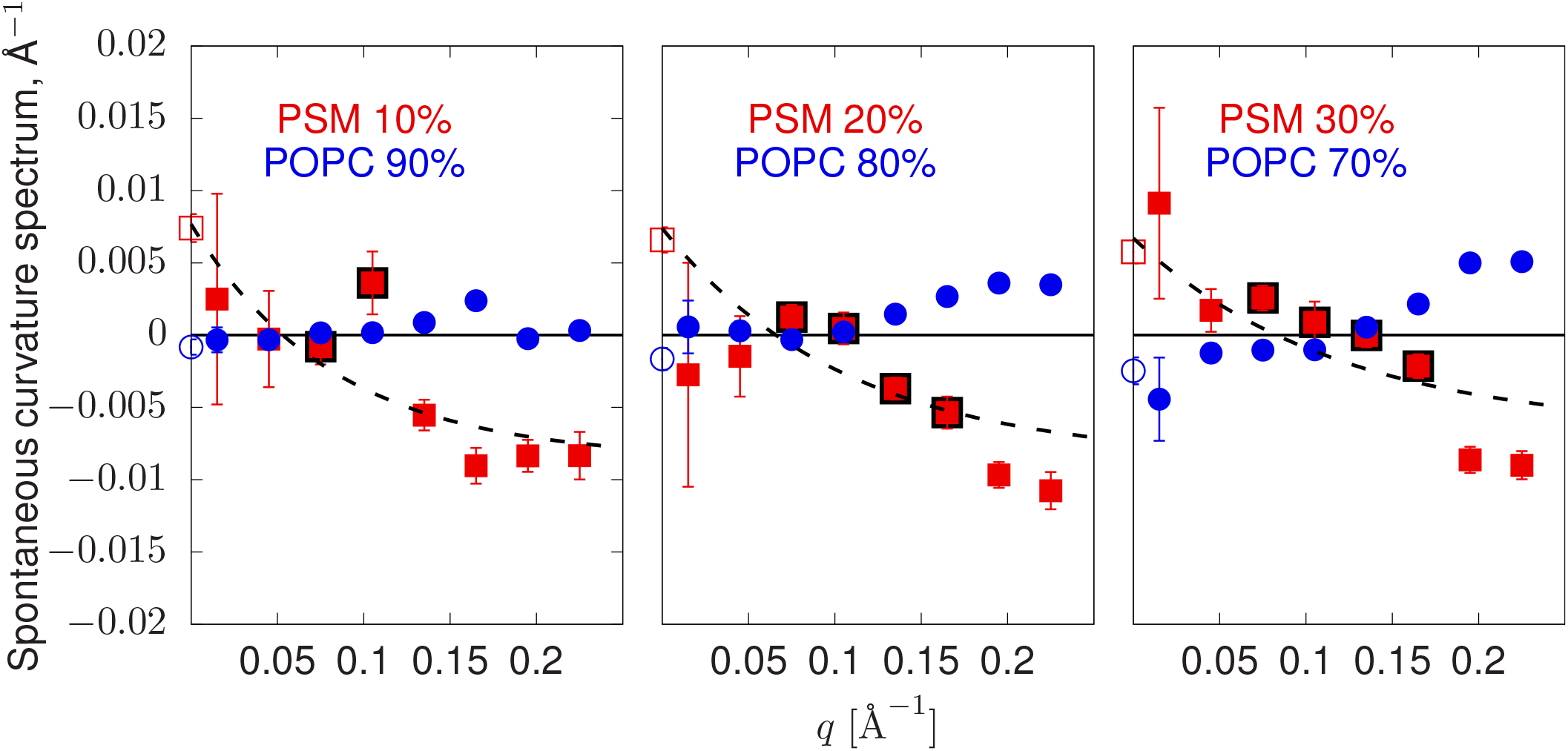
Spontaneous curvature spectra sampled from simulations of PSM in POPC. POPC is shown in blue. PSM is colored red. Open circles at *q* = 0 are computed from the LPP. Each data point is the average over modes within a narrow range of *q*. The dashed line indicates an exponential fit to the spectra as discussed in the text. Points outlined in black in the intermediate range of *q* indicate where saturated lipids appear to have increased positive curvature. Error bars are ± the standard error of the histogram bin mean.

Consider then a lipid analogous to PSM but with a chemically simpler lipid backbone (glycerol). Figure 5 shows the spontaneous curvature spectrum of a 20% DPPC in POPC mixture. Here, a statistically significant peak is indicated between *q* = 0.1 and *q* = 0.15 suggesting a length-scale for the curvature sensitivity of DPPC. These points have been highlighted as for PSM.

**FIG. 5:**
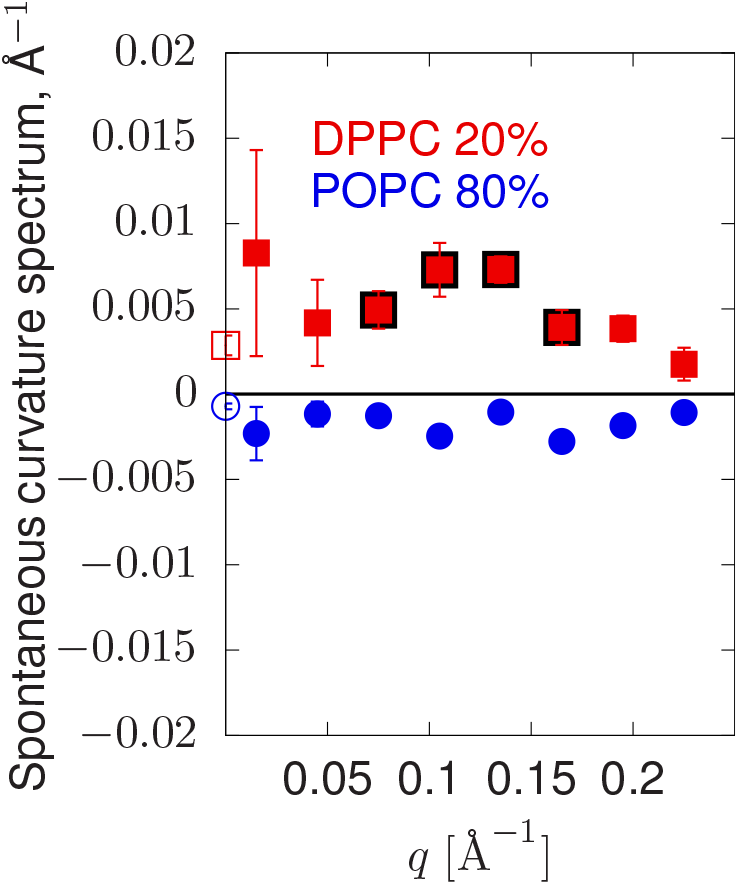
Spontaneous curvature spectra sampled from **DPPC**_20_**POPC**_80_. POPC is shown in blue. DPPC is colored red. The data is a histogram of values. Points outlined in black in the intermediate range of *q* indicate where saturated lipids appear to have increased positive curvature. Error bars are ± the standard error of the histogram bin mean.

Compared with unsaturated lipids, the mechanical extent for these saturated lipids is non-monotonic. Instead, there is a marked positive curvature preference at the nanometer length scale. Considering the development of this feature as PSM concentration is increased, its origins may involve coupling to a background of other saturated lipids present at higher PSM fractions.

A potentially interesting observation at high *q* is that PSM appears to orient according to curvature. In the supporting material, plots of the transverse curvature bias are shown at high *q* for PSM, DPPC, and POPC. The two acyl chains appear to favor substantially different curvature, indicating that the orientation of PSM couples to curvature.

#### C. Non-local mechanical extent supports modulation of lipid phase separation

In 1986, Leibler [60], and later with Andelman [61] applied the Ginzburg-Landau (GL) formalism to characterize how particle-curvature coupling leads to inhomogeneity of the particle distribution and enhancement of surface undulations. The GL formalism is sufficiently flexible to describe line tension physics that gives rise to macroscopic phase separation [62–64]. Lipid composition is represented by the order parameter Φ(*r*) with Fourier weight Φ_**q**_, while the coupling between surface and composition is parameterized with Λ:

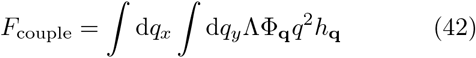

When surface undulations are accounted for, a length-scale *q*\* emerges for the largest variations in Φ_**q**_:

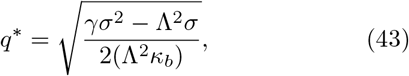

where *σ* is tension and *γ* is the GL parameter penalizing gradients of Φ and is related to line tension [62]. Curvature-compositional coupling can enhance the ability of particles to phase separate by reducing the energetics of variations in Φ, as described in Ref. [24].

This mechanism has been proposed to explain the modulation of the shape of phase-separated lipid domains [23–25, 65], which would otherwise be expected to be circular. For example, stripes emerge in some complex lipid mixtures [22] with a characteristic width that can be tuned. Here the width of the striped phase may be related to 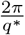. The strength of the emergence of the length-scale *q*\* is set by the relative strength of tension and curvature, e.g., 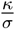. As the magnitude of a thermal undulation is proportional to 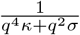, at low *q* tension strongly suppresses membrane undulations relative to high *q*. Thus, a balance is struck for *q*\* at some finite value.

The ME is now introduced. Instead of coupling Φ directly to *h* point-wise, an intermediate function *f*_extent coupled_ is introduced by convolving *w* with Φ:

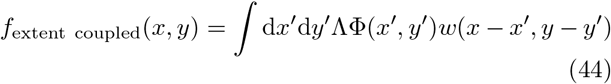

which in Fourier space is simply

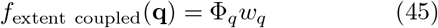

So that Eq. 42 is modified as:

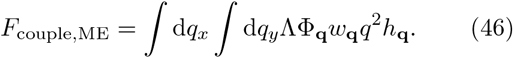

With Eq. 46, new length-scales emerge *regardless of the tension*; they may be set by length-scales intrinsic to *w*, if present. Figure 5 shows a peak in the spontaneous curvature spectrum of DPPC between *q* = 0.1 and *q* = 0.15 Å^−1^ (e.g., a wavelength between 4 and 6 nm). This length-scale is perfectly compatible with domains of ordered DPPC that appear in mixtures with DOPC and cholesterol [66]. Meanwhile, the spectrum for PSM (Figure 4) is somewhat more challenging to interpret but still compelling: there is a statistically significant increase in the spontaneous curvature between *q* = 0.05 and *q* = 0.15, similar to the case as for DPPC. However, PSM appears to have a stronger positive curvature effect at low *q*, at least within the higher PSM background of 20–30%. Note that the spectrum for DPPC and PSM here may be influenced by this very effect—the lipid distribution (i.e. Φ) couples to membrane undulations at a peak value of *q*\*, enhancing the apparent curvature spectrum at that point.

## II. CONCLUSIONS

This work developed the methodology necessary to describe the spatial range of a lipid’s influence on bilayer mechanical properties, which we termed the mechanical extent (ME). The ME influences the position-dependent energy of a lipid. If the ME is highly localized, as we found to be typical of the lipids we modeled here with all-atom molecular dynamics simulation, lipids sense local curvature more strongly than if the effect is distributed over a larger area. Similarly, the ME should influence the chemical potential of lipids as they are trafficked between bilayers; for example, fully localized lipids cannot “balance” the stresses of nearby lipids.

An important step in the method is to control for how to sample the lateral position of a lipid that is composed of many atoms. To do so, we demonstrate how a particular choice, the so-called neutral surface of bending, yields an unbiased position. As a corollary, described in the Appendix, we show how to infer the bending modulus of a lipid from its depth-dependent sampled curvature.

Extracting the undulation-wavelength-dependent average curvature yields what we term the *spontaneous curvature spectrum*. The spectrum is proportional to the ME. Spontaneous curvature spectra for mixtures of varied headgroups and simple unsaturated acyl chains (DOPC, DOPE, and DOPS) demonstrate well-localized extent. While a similar localized curvature effect was found for saturated lipids (PSM and DPPC), they also displayed a marked positive curvature preference at the nanometer length-scale. We predict that this preference for a finite length-scale modulates the shape of lipid-liquid domains, promoting very small domains via curvature coupling.

## III. APPENDIX: COMPUTING THE BENDING MODULUS FROM THE TRANSVERSE CURVATURE BIAS

In Ref. [67] the authors outline how to compute the bending modulus from the undulations of a dynamic molecular simulation without running afoul of apparent fluctuations at higher *q* that are unrelated to curvature energetics. An example of these spurious increases in the undulation spectral intensity are fluctuations of the lipid tilt vector away from the local bilayer normal, as well as protrusions of the lipids above or below their neighbors. The principal result of that work is that by controlling for lipid tilt, the true curvature-mediated undulations can be extracted and the analysis can be extended to higher *q* (shorter wavelength modes amenable to smaller simulations). A similar method was reported by Allolio et al. [68].

Connecting the average curvature sampled by the lipids of a bilayer provides an arguably more convenient route with the same logic. Eq. 16 is based on the correlations of surface curvature and the lipid director. Molecular fluctuations uncorrelated with the surface normal (for example, local tilting) will, on average, not contribute to 〈*c*〉 (*z*). The method is thus to plot the average curvature sampled by a lipid as a function of *z_i_*, with *z_i_* determined by the average height profile of atoms from an approximately planar simulation. The slope of the best fit line to 〈*c*〉 (*z*)_**q**_ is 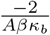.

## ACKNOWLEDGMENTS

This work was supported by the intramural research program of the *Eunice Kennedy Shriver* National Institutes of Child Health and Human Development (NICHD) at the National Institutes of Health. A.H.B. was supported by a Postdoctoral Research Associate (PRAT) fellowship from the National Institute of General Medical Sciences (NIGMS), award number 1Fi2GM137844-01. Simulations were performed on computational resources provided by the NICHD.

