## Supplemental Material for "The spatial extent of a single lipid’s influence on bilayer mechanics"

### Supplemental information for "The spatial extent of a single lipid's influence on bilayer mechanics"

Kayla C. Sapp and Alexander J. Sodt*
*Eunice Kennedy Shriver National Institute of Child Health and Human Development, Bethesda, MD, USA.*

Andrew H. Beaven
*National Institute of General Medical Sciences, Bethesda, MD, USA. and*
*Eunice Kennedy Shriver National Institute of Child Health and Human Development, Bethesda, MD, USA.*

#### $w(q)$ FIT PARAMETERS

Fit parameters for the mechanical extent are shown in Fig. S1. The first column, $k_{\mathrm{local}}\Delta c_0 A_p$, indicates the fraction of the lipid-area-weighted spontaneous curvature is in the entirely local (constant) portion of $w$. The second column, $k_{\mathrm{local}}\Delta c_0 A_p$, is the pre-factor for the exponentially decaying Fourier term in the Fourier representation of $w$. The third column is the spatial decay of that exponential term.

| Lipid | System | $k_{\mathrm{local}}\Delta c_0 A_p$ (Å) | $k_{\mathrm{non-local}}\Delta c_0 A_p$ (Å) | $\alpha$ (Å) |
|---|---|---|---|---|
| DOPC | **PC$_{55}$PS$_{45}$** | 0.0077 | 0.0021 | 1.09967 |
| | **PC$_{25}$PE$_{30}$PS$_{45}$** | -0.00075 | 0.0011 | 0.121298 |
| DOPE | **PE$_{55}$PS$_{45}$** | -0.0056 | -0.0031 | 3.87818 |
| | **PC$_{25}$PE$_{30}$PS$_{45}$** | -0.0134 | — | — |
| DOPS | **PE$_{55}$PS$_{45}$** | 0.0041 | 0.0062 | 2.92603 |
| | **PC$_{55}$PS$_{45}$** | 0.0022 | -0.0069 | 0.950815 |
| | **PC$_{25}$PE$_{30}$PS$_{45}$** | 0.0012 | 0.0039 | 2.40736 |
| PSM | **PSM$_{10}$POPC$_{90}$** | -0.0086 | 0.0162 | 1.9151 |
| | **PSM$_{20}$POPC$_{80}$** | -0.0086 | 0.0160 | 1.497 |
| | **PSM$_{30}$POPC$_{70}$** | -0.0066 | 0.0133 | 1.3457 |
| DPPC | **DPPC$_{20}$POPC$_{80}$** | 0.0032 | — | — |

TABLE S1: Fit parameters using an exponential for $w$. Curvature differences $\Delta c_0$ depend on the total composition of the bilayer and so vary for a single lipid between simulations.

#### ALIGNMENT OF PSM ORIENTATION TO CURVATURE IS INDICATED BY THE TRANSVERSE CURVATURE BIAS AT HIGH $q$.

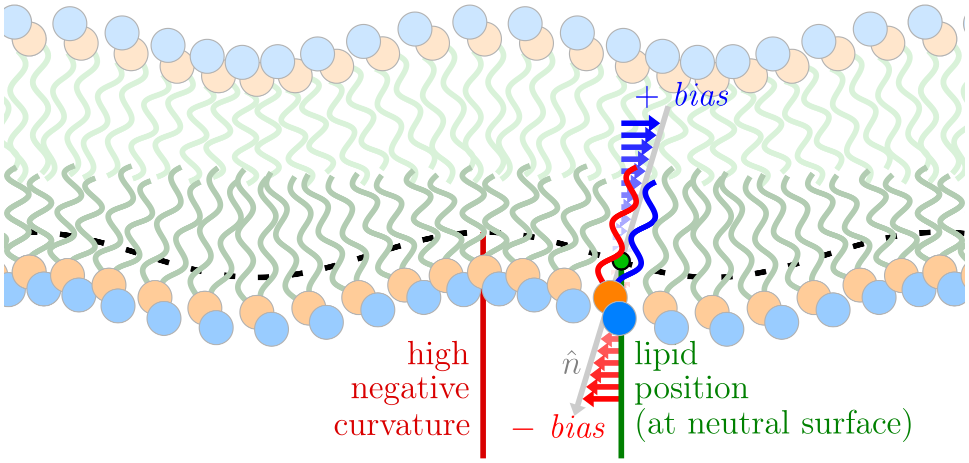


FIG. S1: A cartoon showing how curvature/orientational coupling leads to a bias between acyl chains.

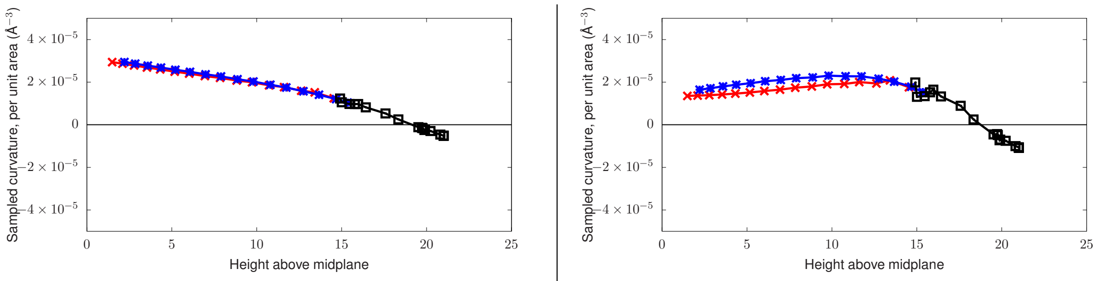


(a) $0.06 < q < 0.075$ (b) $0.18 < q < 0.195$

FIG. S4: The transverse curvature bias of a DPPC lipid in a 20% DPPC/80% POPC simulation. The *sn*-1 chain is colored red, while the *sn*-2 chain is colored blue.

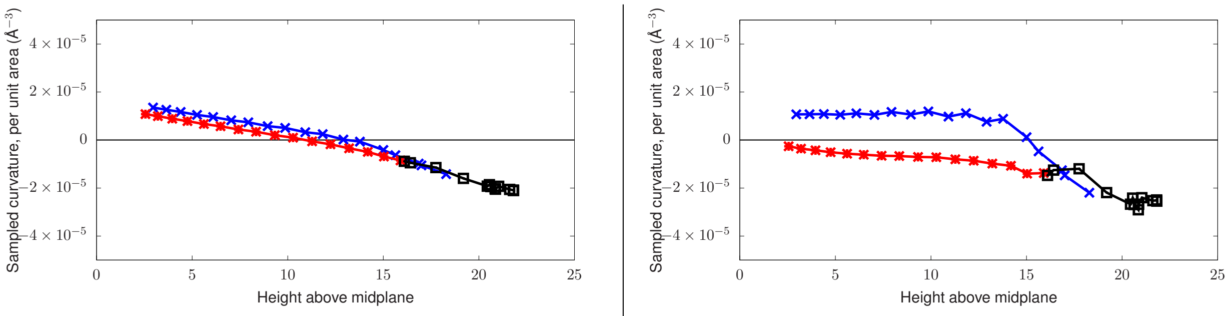


(a) $0.06 < q < 0.075$ (b) $0.18 < q < 0.195$

FIG. S2: Transverse curvature bias of PSM in a 10% PSM/90% POPC simulation. The coloring for the imagined sphingosine acyl chain is blue, while the fatty-acid acyl chain is shown in red.

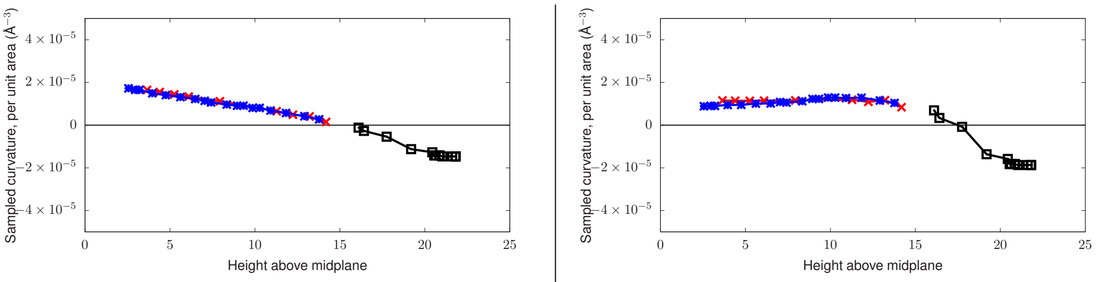


(a) $0.06 < q < 0.075$ (b) $0.18 < q < 0.195$

FIG. S3: The transverse curvature bias of a POPC lipid in a 10% PSM/90% POPC simulation. The *sn*-1 chain is colored red, while the *sn*-2 chain is colored blue.

Figures S2a and S2b shows the apparent curvature experienced by a PSM lipid, measured by various atoms throughout the lipid. The striking feature in Fig. S2b is that one acyl chain of PSM, the sphingosine chain, tends to be oriented toward positive curvature, while the fatty-acid derived chain tends to be oriented toward negative curvature.

The feature is either absent or dramatically diminished for POPC and DPPC; see Figs. S3a, S3b, S4a, and S4b.

Chemical schematics of PSM, POPC and DPPC are shown in Fig. S5.

* Electronic address:

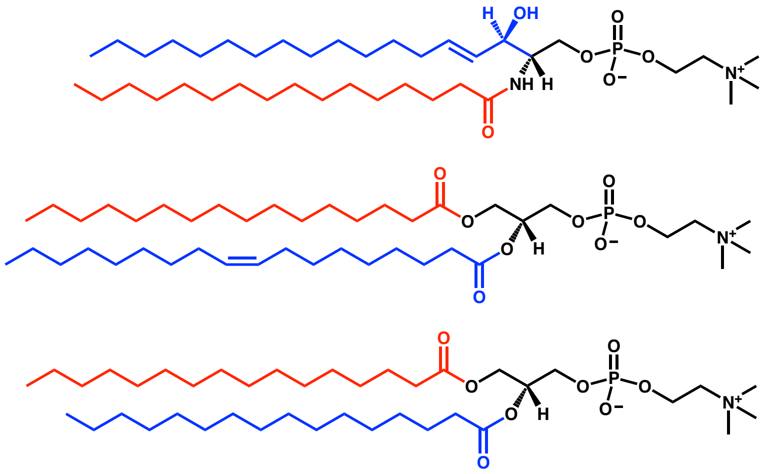

PSM

POPC

DPPC

FIG. S5: Schematics of the lipids with transverse curvature bias shown in this Supplemental Material.
